# Differential haplotype expression in class I MHC genes during SARS-CoV-2 infection of human lung cell lines

**DOI:** 10.1101/2022.11.20.517193

**Authors:** Ronaldo da Silva Francisco Junior, Jairo R. Temerozo, Cristina dos Santos Ferreira, Yasmmin Martins, Thiago Moreno L. Souza, Enrique Medina-Acosta, Ana Tereza Ribeiro de Vasconcelos

**Author notes:** **Corresponding author:** Ana Tereza Ribeiro de Vasconcelos,.

## Abstract

Cell entry of SARS-CoV-2 causes genome-wide disruption of the transcriptional profiles of genes and biological pathways involved in the pathogenesis of COVID-19. Expression allelic imbalance is characterized by a deviation from the Mendelian expected 1:1 expression ratio and is an important source of allele-specific heterogeneity. Expression allelic imbalance can be measured by allele-specific expression analysis (ASE) across heterozygous informative expressed single nucleotide variants (eSNVs). ASE reflects many regulatory biological phenomena that can be assessed by combining genome and transcriptome information. ASE contributes to the interindividual variability associated with disease. We aim to estimate the transcriptome-wide impact of SARS-CoV-2 infection by analyzing eSNVs. We compared ASE profiles in the human lung cell lines Calu-3, A459, and H522 before and after infection with SARS-CoV-2 using RNA-Seq experiments. We identified 34 differential ASE (DASE) sites in 13 genes (*HLA-A, HLA-B, HLA-C, BRD2, EHD2, GFM2, GSPT1, HAVCR1, MAT2A, NQO2, SUPT6H, TNFRSF11A, UMPS*), all of which are enriched in protein binding functions and play a role in COVID-19. Most DASE sites were assigned to the MHC class I locus and were predominantly upregulated upon infection. DASE sites in the MHC class I locus also occur in iPSC-derived airway epithelium basal cells infected with SARS-CoV-2. Using an RNA-Seq haplotype reconstruction approach, we found DASE sites and adjacent eSNVs in phase (i.e., predicted on the same DNA strand), demonstrating differential haplotype expression upon infection. We found a bias towards the expression of the HLA alleles with a higher binding affinity to SARS-CoV-2 epitopes. Independent of gene expression compensation, SARS-CoV-2 infection of human lung cell lines induces transcriptional allelic switching at the MHC loci. This suggests a response mechanism to SARS-CoV-2 infection that swaps HLA alleles with poor epitope binding affinity, an expectation supported by publicly available proteome data.

## Introduction

The coronavirus disease 2019 (COVID-19) pandemic significantly continues to burden public health response and management, with over 631 million infected people and over 6.5 million cumulative deaths worldwide (https://covid19.who.int/). The severe acute respiratory syndrome coronavirus 2 (SARS-CoV-2) infection causes asymptomatic to life-threatening pulmonary illness with multiorgan dysfunction (1,2). About 0.1 to 0.9% of infected people evolve fatal disease outcomes (3).

Epidemiological studies showed that advanced age, male biological sex, and comorbidities are major risk factors for life-threatening COVID-19 (4). Respiratory tract epithelial cells and pneumocytes are the first target cells of SARS-CoV-2. The virus enters cells through the binding of the Spike protein to the host angiotensin-converting enzyme 2 (ACE2) membrane receptor (5). The kinetics of the SARS-CoV-2 replicative cycle during the acute phase of infection can lead to endothelial barrier disruption, dysfunctional alveolar-capillary oxygen transmission, and impairment in oxygen diffusion capacity (6). These phenotypes are characteristic of acute respiratory distress syndrome (ARDS) and usually demand oxygen support.

A hallmark of severe COVID-19 is the overactivation of the inflammatory response through maladaptive proinflammatory cytokine production by transendothelial leukocyte migration. The cytokine storm causes local cell damage in the alveoli and systemic inflammation. Excessive inflammation, hypoxia, immobilization, and diffuse intravascular coagulation are not uncommonly observed in COVID-19 patients. Those conditions may predispose to both venous and arterial thromboembolism, ischemic stroke, and myocardial infarction, which are life-threatening complications (7). Also, SARS-CoV-2 interferes with the way antigens are presented, how alveolar macrophages work, and how type I interferon works (8,9).

Understanding the perturbations associated with SARS-CoV-2 infection in the respiratory tract cells is challenging because of the difficulty in obtaining relevant biological samples from affected subjects. To do this, *in vitro* culture models permissive to SARS-CoV-2 infection are used to investigate the underlying mechanisms of infection and disease pathology. Calu-3 and A549 are the most amenable human lung-derived cell line models available (10,11). Even though both cells are epithelial and come from adult lung non-small cell adenocarcinoma, Calu-3 is highly permissive to SARS-CoV-2 infection and replication in an ACE2-dependent way, while A549 is not permissive to SARS-CoV-2 due to its low expression of the ACE2 receptor (12). Notably, the exogenous expression of ACE2 in A549 renders a chemokine signature similar to Calu-3 cells (12). ACE2 receptor-independent models, such as the H522 lung adenocarcinoma cell line, showed that viral infection uses an alternative receptor and depends on surface heparan sulfates (13). In addition, airway epithelium basal cells (iBCs) experimentally derived from induced pluripotent stem cells (iPSCs) also reproduced the transcriptome profile of the primary human airway epithelial cells and other airway cell types (14). Comprehensive transcriptome studies with these cell lines showed genome-wide activation of genes related to type I and III IFN production, chemokine expression, NLRP3 inflammasome, metabolic hormone process, and low-density granulocyte (LDG) gene signature (12,15–17). Nevertheless, the impact of SARS-CoV-2 infection on allele-specific expression has not been fully explored.

Genome-wide association studies (GWAS) found that common SNVs at 17 different loci were linked to severe COVID-19 outcomes (18–20). Loss-of-function rare SNVs in genes related to inborn errors of type I IFN immunity were found in at least 3.5% of patients with pneumonia (3,21). Variants in the Human Leukocyte Antigen (HLA) locus appear to play a role in asymptomatic and mild diseases. The highly variable HLA locus codes for proteins that activate T-cells and help the immune system fight off different pathogens. Class I and II HLA molecules present antigens to CD8+ and CD4+ T-cells, respectively. In couples who were discordant for COVID-19, HLA-A variants were associated with highly exposed, asymptomatic, seronegative women (22). In resilient super elders (i.e., infected individuals aged over 90 years presenting with mild or no symptoms), an increased frequency of missense variants in the *MUC22* gene was found (23).

Most of the markers found by GWAS are single-nucleotide variations (SNVs) at noncoding sites that often act as cis-regulatory variants. Expression quantitative trait loci (eQTL) analysis (24,25) is often used to identify causal regulatory variants from GWAS, which also requires many samples, despite being deeply influenced by interindividual differences (26). SARS-CoV-2 infection is known to promote imbalances in the expression of genetic variants across the human genome (27). But it is still not clear what their functional effects are because figuring out the links between genotype and phenotype in people with different genetic backgrounds requires analyzing a large number of transcriptomes. For these reasons, allele-specific expression (ASE) has become the most effective assay for quantification of genetic variant expression (28).

ASE analysis measures the steady-state imbalance in the transcription of the two parental alleles at heterozygous sites of the diploid genome (29). Each genetic variation is expected to show a 1:1 allelic expression ratio. The departure from this assumption captures a dynamic regulation of biological processes related to the effects of cis-regulatory variants, genomic imprinting, X chromosome inactivation (XCI), A-to-I(G) RNA editing, nonsense-mediated decay, random monoallelic expression, or allelic exclusion (30). ASE analysis also helps the identification of gene-by-environment (GxE) interactions, highlighting the environment’s contributions to modulating the genetic effects of relevant complex traits (31). Unlike GWAS and eQTL analyses, ASE analysis quantifies the allelic effects within the same individual, by controlling the effects of genetic background and environmental changes in replicate samples (26). Comparisons across allelic expression profiles can highlight genes potentially involved in mechanisms associated with the disease. For example, Goovaerts et al., found that the parent-of-origin-dependent monoallelic expression of imprinted genes is deregulated in breast cancer (32). Pervasive perturbations in ASE sites were found in monozygotic twins discordant for Down syndrome, suggesting genome-wide dysregulation in cells with extranumerary chromosome 21 (30). Here, we describe a new way to use RNA-Seq experiments on human cell lines infected with SARS-CoV-2 to find allele-specific changes that are important for COVID-19 disease.

## Materials and methods

### Biological data and sample information

We chose transcriptome studies from publicly available bulk RNA-Seq data of SARS-CoV-2 infected lung human cell lines on the Sequence Read Archive platform (Table S1). Only experiments comparing mock-treated and SARS-CoV-2 infected cells with two or more replicates per condition were selected. We included three different lung cell lines in our analysis: Calu-3, A549, and H522. These cell lines originated from the lung adenocarcinoma epithelium of Caucasian adult male subjects. Both A549 and H522 are ACE2-negative models supporting SARS-CoV-2 replication via independent entries. In our analysis, we also used A549 with an exogenous expression of ACE2. In the study by Blanco Melo et al., 2020 (GEO BioProject PRJNA615032), we selected four experiments using the A549 cell line and one from Calu-3 (12). In the study by Wyler et al. (GEO BioProject PRJNA625518), we included a longitudinal experiment of RNA-Seq in Calu-3 cells at three different time points (16). We also used RNA-Seq data of Calu-3 cells from the study by Kim D et al., 2021 (GEO BioProject PRJNA661467) at eight different time points (17). The H522 experiments were retrieved from the study conducted by Puray-Chavez et al. (GEO BioProject PRJNA686659), which compares the transcriptional profile for four ratios of the multiplicity of infection (MOI) at six-time points (13). We also use data from whole-exome sequencing (WES) data for the cell lines listed above (Table S1) to figure out the zygotic profile of each RNA-Seq variant. Lastly, we used RNA-Seq data of airway epithelium basal cells (iBCs) made from induced pluripotent stem cells (iPSCs) (GEO BioProject PRJNA805095) to confirm what we found in the three models we used in our analysis. The human airway epithelium cells were differentiated from BU3 NGPT and 1,566 iPSC lines (14).

### Data processing and identification of differential allele-specific expression sites

We extracted the *fasta* files of each replicate using the *fastq-dump* function from the *sra-toolkit* (https://github.com/ncbi/sra-tools). Bioinformatic analysis was conducted separately for each replicate. Allelic imbalance analysis at expressed SNVs (eSNVs) sites was performed using *PipASE*, a pipeline to identify ASE sites in transcriptome data (30). We first examined the sequencing quality parameters for each *fastq* file using *fastqc* (https://www.bioinformatics.babraham.ac.uk/projects/fastqc/). Next, bad-formed reads were removed using *Trimmomatic* (33). We aligned the filtered reads to the human GRCh38 reference genome assembly with STAR v3.7 software (34). Mapped sequences were further post-processed using *SAMtools* to sort, index, and select reads based on mapping quality parameters (MAPQ ≥ 30) in *BAM* files (35). Then, we masked duplicate reads and performed variant calling in RNA-seq data using *MarkedDuplicates* and *HaplotypeCaller* from GATK v4.1, respectively (36,37). We used *ASEReadCounter* to determine the counts for reference and alternative alleles in each position (29). The genomic information for each variant was annotated using the Ensembl Variant Effect Predictor (https://www.ensembl.org/Tools/VEP).

To estimate the impact of SARS-CoV-2 infection on the differential expression of genetic variants across the human genome, we calculated the reference allele ratio (ref ratio) in each replicate using the following equation: ref ratio = (# of reads with the reference allele) / (# of reads with the reference allele + # of reads with the alternative allele). For differential ASE analysis, we required coverage of at least ten reads per variant site and the occurrence of each site in at least two replicates in each assay condition. We used a binomial model from the stats package R (38) to do differential ASE analysis at each eSNV site. Adjusted P-values for multiple comparisons were performed using the *p.adjust* function in R with the Benjamini & Hochberg method. To estimate the magnitude of the expression changes, we calculated the log2 fold change of the ASE (LogASE) for each site using *DESeq2* (39) according to the framework available by Love (2017). Positive LogASE values represent the increase of the alternative allele over the reference. In contrast, negative values represent ASE sites that exhibited a preferential expression of the reference allele after infection. Only the SNVs that exhibited FDR < 0.1 and −0.95 < LogASE > 0.95 were considered differentially expressed across the conditions. We used the R package *clusterProfiler* to perform functional enrichment analysis on the set of genes that displayed differential allele expression in our study (40). Annotations were made for Gene Ontology (GO) terms in three different areas: molecular function (MF), biological process (BP), and cellular component (CC). We performed a GO over-representation test, keeping only enriched terms that showed *p.adjust* < 5%. To further investigate the main metabolic pathways enriched for the genes containing ASE sites, we also conducted a KEGG over-representation analysis using *clusterProfiler*. Similar analyses were also performed using *ReactomePA* in R (41).

### Detection of chromosomal aberrations and haplotype inference using allelic imbalance from RNA-seq dataset

As the tumoral cell lines used in this study are hypotriploid (42), we conducted a chromosomal aberration analysis in RNA-Seq data using eSNP-Karyotyping (43). We sought to compare the karyotype of mock-treated and SARS-CoV-2-infected cells to determine whether the allelic imbalance was either generated by chromosomal differences between both samples or associated with the infection. Thus, *BAM* files from different replicates within the same condition were merged with *SAMtools* (35) and edited using *AddOrReplaceReadGroups* from Picard (https://broadinstitute.github.io/picard/) to assign a single new read-group for all reads in the *BAM* file. The *BAM* file generated by this step was indexed with the SAMtools index, followed by a second variant calling with HaplotypeCaller from GATK v4.1. We filtered out eSNVs with low coverage (below 20 reads) and low minor allele frequency (lower than 0.2). Using a window of 151 eSNVs, we estimate the moving medians of the major to minor allele ratios across the genomic coordinates. eSNP-Karyotyping also shows FDR-corrected P values for regions significantly altered within each sample. Combined *BAM* and *VCF* files were also used to phase eSNVs within haplotype blocks. We used a Bayesian haplotype reconstruction framework from HapTree-X to assess phased haplotype blocks from the allelic imbalance observed in RNA-Seq data (44). We passed the human *GTF* file from the Ensembl GRCh38.105 version via the -g parameter to improve the phasing quality.

### Sequence-based HLA typing using RNA-Seq data

After the haplotype reconstruction approach, we conducted HLA allele identification directly from RNA-Seq reads in each sequence. First, RNA-Seq reads in *fastq* format were mapped to human chromosome 6 (GRCh38) using *bowtie2* (45). The mapped sequences were assembled into 200 bp contigs using the *TASR* tool (46) and aligned to HLA reference sequences by using the NCBI BLAST+ 2.13.0 package (https://blast.ncbi.nlm.nih.gov/Blast.cgi). The following alignment parameters were used: -b 5 -v 5. The HLA reference sequences of classes I and II genotypes were retrieved in *fasta* format from the IMGT/HLA database. After alignment, the selected sequences were used to predict HLA alleles in the HLAminer tool with the default parameters (47). Next, the definition of HLA alleles for each sample was based on the intersection of alleles present across the different replicates of the experiments. Finally, we queried DASE sites and co-localized eSNVs affected in samples predicted to be heterozygous to verify the HLA allele preferentially expressed during SARS-CoV-2 infection.

## Results

### Allelic expression of eSNVs in MHC class I locus is preferentially impacted in lung epithelial cell lines during SARS-CoV-2 infection

We compared the allelic expression profiles of eSNVs in bulk RNA-Seq data from Calu-3, A549, and H522 lung cell lines before and after SARS-CoV-2 infection. We interrogated 6,884 heterozygous eSNVs detected across the mock-treated and SARS-CoV-2-infected comparisons, with coverage ≥ 35 reads at each site. Thirty-four eSNVs displayed differential allele-specific expression (DASE) after viral infection (Figure 1; Table S2). These sites were heterozygous in the WES data of their respective cell lines. The ACE2-dependent model, Calu-3 (n = 23/2,850), harbored 68% of all DASE sites. We also noticed seven eSNVs significantly altered in A549 with exogenous expression of ACE2 (n = 7/4,094). The ACE2-independent models of H522 and A549 showed the smallest DASE sites with four (n = 3/672) and two eSNVs (n = 2/872), respectively (Figure 1). The read depth at DASE sites was 2.5-fold greater than the coverage across all positions.

**Figure 1.**
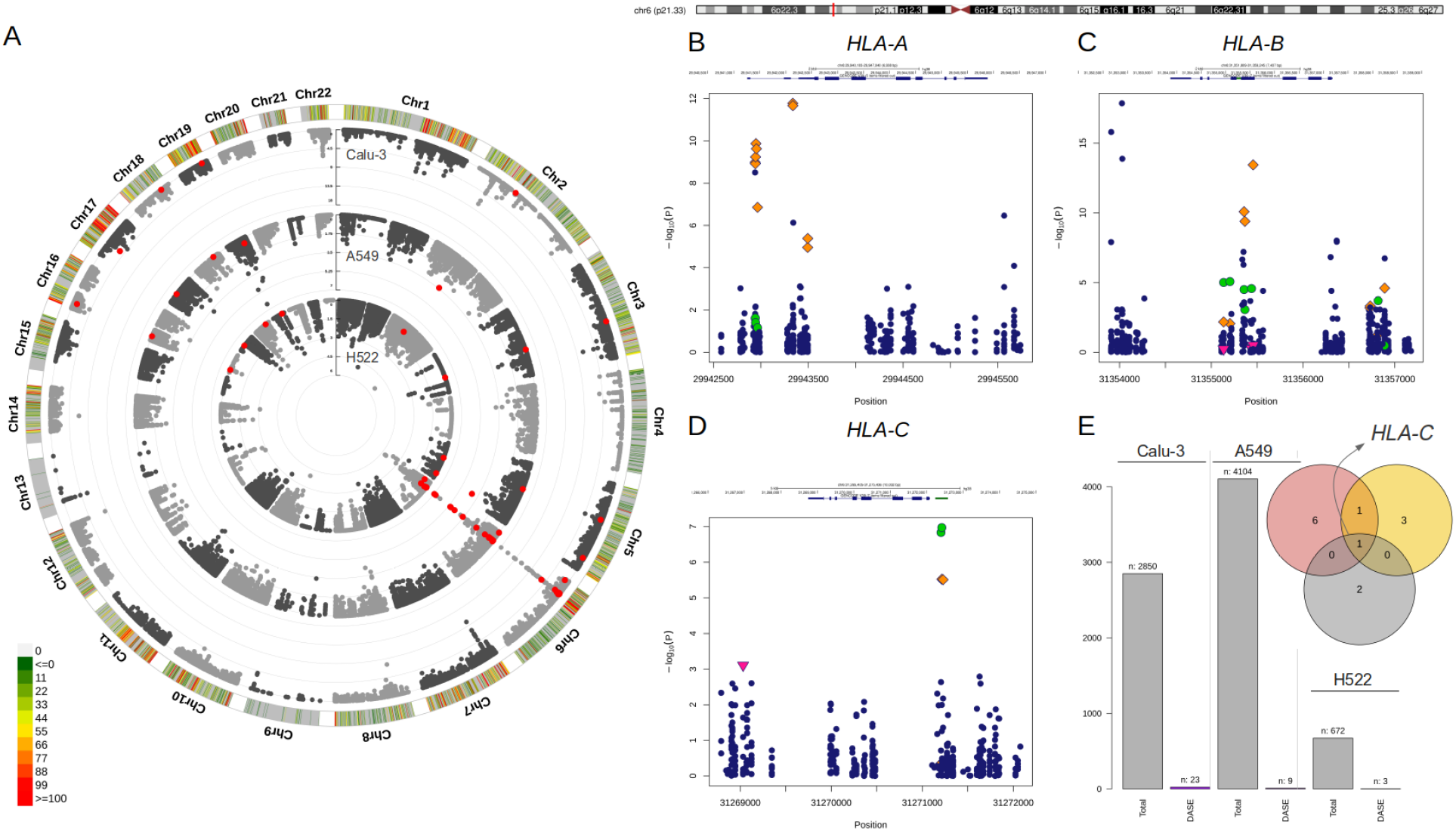
Differential allele-specific expression sites across the single-nucleotide variants identified in Calu-3, A549, and H522 lung cell lines. A) Circular Manhattan plot of the chromosomal distribution of eSNVs tested using a binomial approach. The densities of eSNVs per chromosome in the Calu-3, A549, and H522 cell lines are depicted inward. Red points represent DASE sites with FDR < 10%. B-D) Regional plot of classical MHC class I genes with the orange diamond showing the DASE sites in Calu-3, green circles for A549, and H522 represented by the pink triangle point down. E) The total number of eSNV sites tested in each lung cell line, followed by the number of DASE sites found. The intersection between the genes harboring DASE sites in the three cell lines is depicted in the Venn diagram. *HLA-C* was the only gene that showed DASE sites in all lines. Nevertheless, *HLA-B* was also shared between Calu-3 (red circle) and A549 (yellow circle).

Nineteen DASE sites were mapped to coding regions, with 56% being missense and 41% being synonymous variants. Only one eSNV mapped to the *HLA-C* 3’ UTR. Furthermore, DASE sites are in 13 autosomal genes located on eight chromosomes (Table S2), with most eSNVs on chromosome 6. The major histocompatibility complex (MHC) locus harbored 24 (70%) of the DASE sites (Figure 1). *HLA-B* (n = 10) and *HLA-A* (n = 10) carried the highest number of affected variations followed by *HLA-C* (n = 4). Only one eSNV changed significantly in each of the other ten genes (*BRD2, EHD2, GFM2, GSPT1, HAVCR1, MAT2A, NQO2, SUPT6H, TNFRSF11A, and UMPS). HLA-C* harbored DASE sites in all lung cell lines included in this study (Figure 1). Both Calu-3 and A549 also shared DASE sites in the *HLA-B* gene. No significant association was observed between the number of DASE sites from the different multiplicity of infection (MOI) ratios and hours post-infection (hpi), suggesting that the mechanisms underlying the differential expression of some alleles may be independent of these variables.

Gene ontology (GO) over-representation analysis revealed that upregulated genes are mainly involved in antigen processing and presentation of endogenous peptides via MHC class I (GO:0019885), cell killing (GO:0001906), and regulation of leukocyte mediated cytotoxicity (GO:0001910). We observed an association between *HLA-A* and *HLA-B* with IFN-γ (GO:0032609) and interleukin-12 production (GO:0032615). *GFM2* and *GSPT1* were associated with the biological process of translational termination (GO:0006415). We also noticed an enrichment of the guanyl ribonucleotide binding (GO:0032561) molecular functions linked to *EHD2, GFM2*, and *GSPT1. TNFRSF11A* showed significant over-representation in the tumor necrosis factor-activated receptor (GO:0005031) and death receptor (GO:0005035) activities. *HAVCR1* displayed virus receptor activity (GO:0001618), whereas *NQO2* had a function of chloride ion binding (GO:0031404).

### The expression profiles of the genes harboring DASE sites distinguish genetic regulatory mechanisms triggered by infection

The allelic imbalance observed at DASE sites could result from the differential gene expression (DGE) induced by SARS-CoV-2. So, we compared the LogASE values to the log_2_-fold change (LogFC) of the significant DGE (Figure 2A). We found that 23 DASE sites were linked to increased expression of the *HLA-A, HLA-B*, and *HLA-C* genes at 24 hpi in Calu-3, A459, and H522 cell lines (Table S2). This showed that HLA expression was increased in a way that was specific to each chromosomal copy. Such differentiation was detected across the seven experiments included in our study. For 14 eSNVs in *HLA-A* (n = 7), *HLA-B* (n = 6), and *HLA-C* (n = 1), upregulation was seen in DASE sites where the reference allele was more likely to be expressed (Figure 2B). Ten eSNVs in the upregulated group showed that the alternative allele was more often expressed upon SARS-CoV-2 infection (Figure 2B). In Calu-3 cells, the rs713031 in the *HLA-B* gene showed random allele expression over time, with an allelic imbalance towards the alternative allele at 24 hpi with MOI = 10 and switching to the reference allele at 48 hpi with a 0.1 MOI. For both experiments, an increased transcriptional level was detected for the gene. Such an allele switch may represent a random allelic imbalance expression once both comparisons where this eSNV was detected originated from cells with the same genotype.

**Figure 2.**
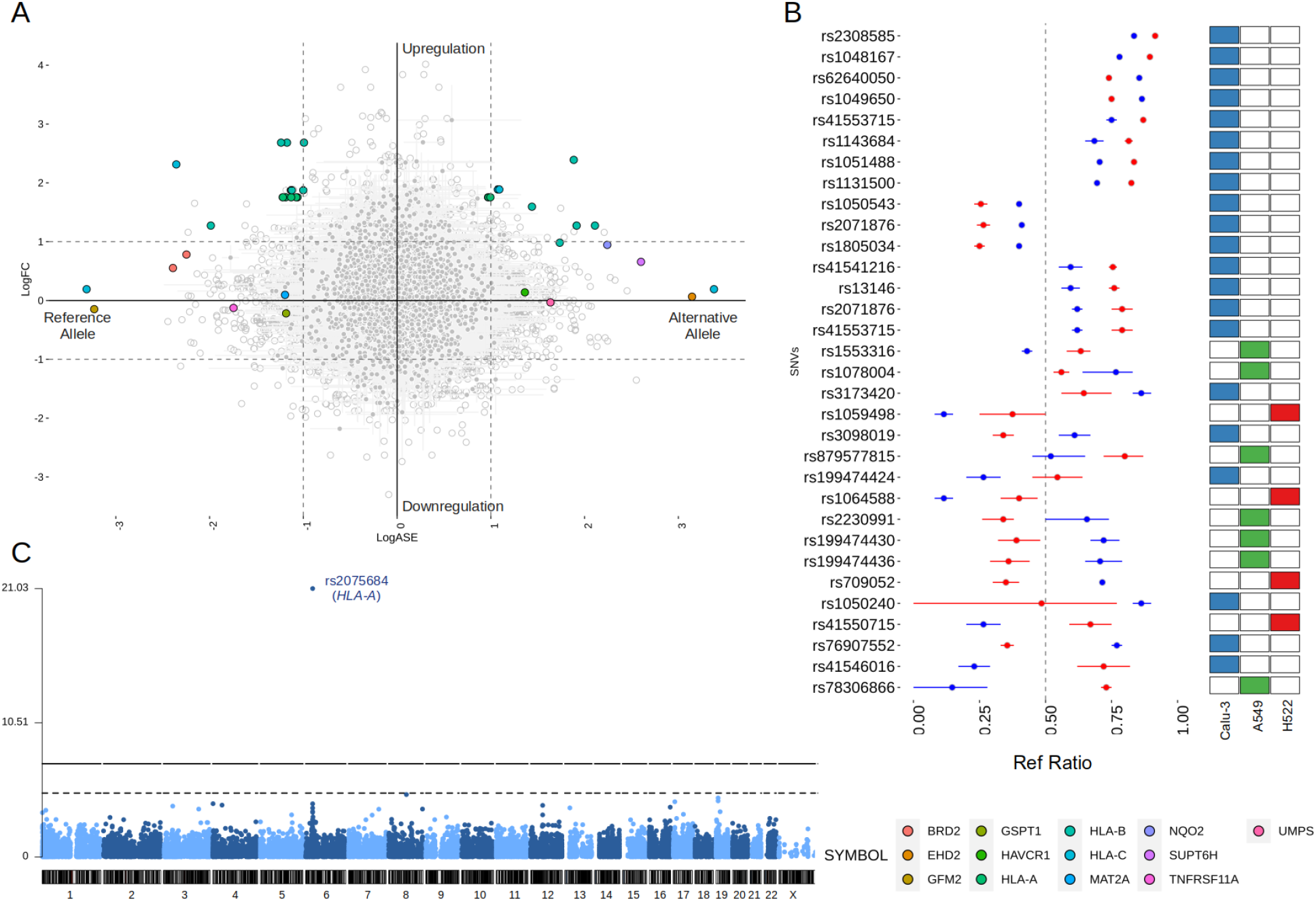
Comparison of the LogASE and LogFC from differential gene expression (DGE). A) Plot showing the LogASE values for DASE sites on the x-axis and LogFC from the DGE comparing the infected with mock-treated cells. Positive LogASE values represent the increase of the alternative allele over the reference. In contrast, negative values represent ASE sites that exhibited a preferential expression of the reference allele after infection. Colored circles show the genes where each DASE site is mapped, whereas gray circles show the eSNVs that met the requirement of FDR > 10% in the DASE analysis. B) Comparison between the Ref Ratio values of SARS-CoV-2 infected and mock-treated cells. The plot shows the ref ratio values on the x-axis and DASE sites on the y-axis. Red and blue circles represent the mean of Ref Ratio values among the replicates of SARS-CoV-2 infected and mock-treated cells, respectively. The interval bars denote the range between the min and max values found across the replicates. Ref Ratio values > 0.5 represent the preferential expression of the reference allele, whereas values < 0.5 show the bias towards the alternative allele. The heatmap at the right highlights the cell line where the DASE sites were found. C) Manhattan plot showing the DASE sites found in iPSC-derived airway epithelium basal cells (iBCs).

Twelve genes with 13 DASE sites showed compensated expression, which means that the virus did deregulate their expression level. At six eSNVs, the reference allele was expressed more than the alternative allele, but at seven DASE sites, the alternative allele was expressed more. No shift in the allele expression of the same eSNV across different experiments was observed in this group of genes. For the rs2071876 in *BRD2*, we identified a consistent expression of the reference allele in the H522 cell line at 72 and 96 hpi (MOI = 0.06).

Furthermore, *HLA-B* and *HLA-C* also displayed compensated gene expression at 12 hpi despite being upregulated at 36 hpi in the same Calu-3 cell (MOI = 10). Though the gene expression changed, for rs41553715 in *HLA-B*, the expression of the alternative allele was increased in both scenarios. Interestingly, the reference allele was preferentially expressed during upregulation of the gene at 48 hpi in Calu-3 (MOI = 0.1) for the same genetic variant, suggesting biased allele expression or a parental-dependent effect. Similar results were found when comparing different *HLA-C* cell lines; for the rs41550715, the alternative allele was preferential regardless of gene expression compensation in A549 or upregulation in Calu-3 (Table S3).

### *HLA-A* gene is also altered in iPSC-derived airway epithelium basal cells

Next, we aimed to verify the expression profiles of genetic variants across alternative cell lines to determine the extension of the DASE events. We then performed the ASE analysis on a dataset of airway epithelium basal cells derived from iPSC lines (iBCs). The iBCs originated from two independent precursors (iBCs-1566 and iBCs-BU3 NGPT). Unlike lung-derived cell lines, we could not retrieve WES data from both cells. Therefore, the genetic variants identified were considered theoretically heterozygous. We interrogated 26,420 sites, including 14,909 from iBCs-1566 and 16,338 from iBCs-BU3 NGPT. The SNV rs2075684-T-A located in the *HLA-A* gene was found to be differentially expressed during viral infection in iBCs-1566 cells after 24 hpi (Figure 2C; Table S3). After infection, the reference T allele was seen to be more active than the alternative A allele. This variation changes phenylalanintoby tyrosine at position 33 (Phe33Tyr) of the HLA-A protein. However, the alleles carrying phenylalanine codons seem to be preferentially expressed. The overall minor allele frequency of rs2075684 was 0.14 in GnomAD. Allele A has a MAF greater than 0.3 in South and East Asia populations. During infection, the expression profiles of two other HLA-A variants, rs45585732 and rs1655894, which are close to each other, were changed. In iBCs (BU3 NGPT), we detected two DASE sites (rs2269350-G-A and rs11724369-G-A) in the *RPSA* and *UVSSA* genes after 72 hpi (Table S3). Both genetic variants had a synonymous effect on their proteins. The expression of the reference alleles went down, and then the expression of allele A went up. The alternative allele has a relatively elevated frequency across the populations in GnomAD (MAF = 0.26 and 0.29, respectively). We also identified expression perturbations across five neighboring SNVs (rs2276903, rs28614045, rs9996817, rs9685761, and rs6838561) in the *UVSSA* gene. All three genes highlighted in iBCs displayed a compensated gene expression profile in the experiments.

### DASE sites are not related to chromosomal aberrations differences between mock-treated and SARS-CoV-2 infected samples

Having identified DASE sites across lung-derived and airway basal epithelial cell lines, we asked whether these allele biases were caused by genomic instability or viral infection. We wished to rule out possible karyotype differences as the primary source of ASE since Calu-3, A549, and H522 are hypotriploid (42). We confirmed chromosomal aberrations in all cell lines analyzed using eSNP karyotyping and WES karyotyping (Figure 3). Though the eSNP-Karyotyping revealed a dynamic pattern in the RNA-Seq data of Calu-3, no significant karyotype alterations were detected at the DASE sites (Figure 3). For A459, the nine DASE sites identified are mapped at chromosomes 2, 5, 6, and 19, of which six SNVs target the MHC class I locus (Figure 3). At the genomic level, we detected significant alterations in chromosomes 17 and 20. Both aberrations were also present at the transcriptional level reported by eSNP-Karyotyping analysis in all A549 experiments. The pattern was consistent when both conditions were compared. Karyotyping with WES or RNA-Seq data in H522 cells suggested the presence of a structural aberration across the MHC locus (Figure 3, panels E and F). Infected cells have a DASE site in the HLA-C gene. This suggests that the observed allelic shift is related to SARS-CoV-2. We could not retrieve WES data from the iBCs lines used in our study. Despite this, the allelic ratios from RNA-Seq data were consistent in both IBC cell lines, implying that no chromosomal aberrations were present (Figure 3). Thus, the DASE sites are not likely to be caused by the alterations in the karyotype of the mock-treated and infected samples.

**Figure 3.**
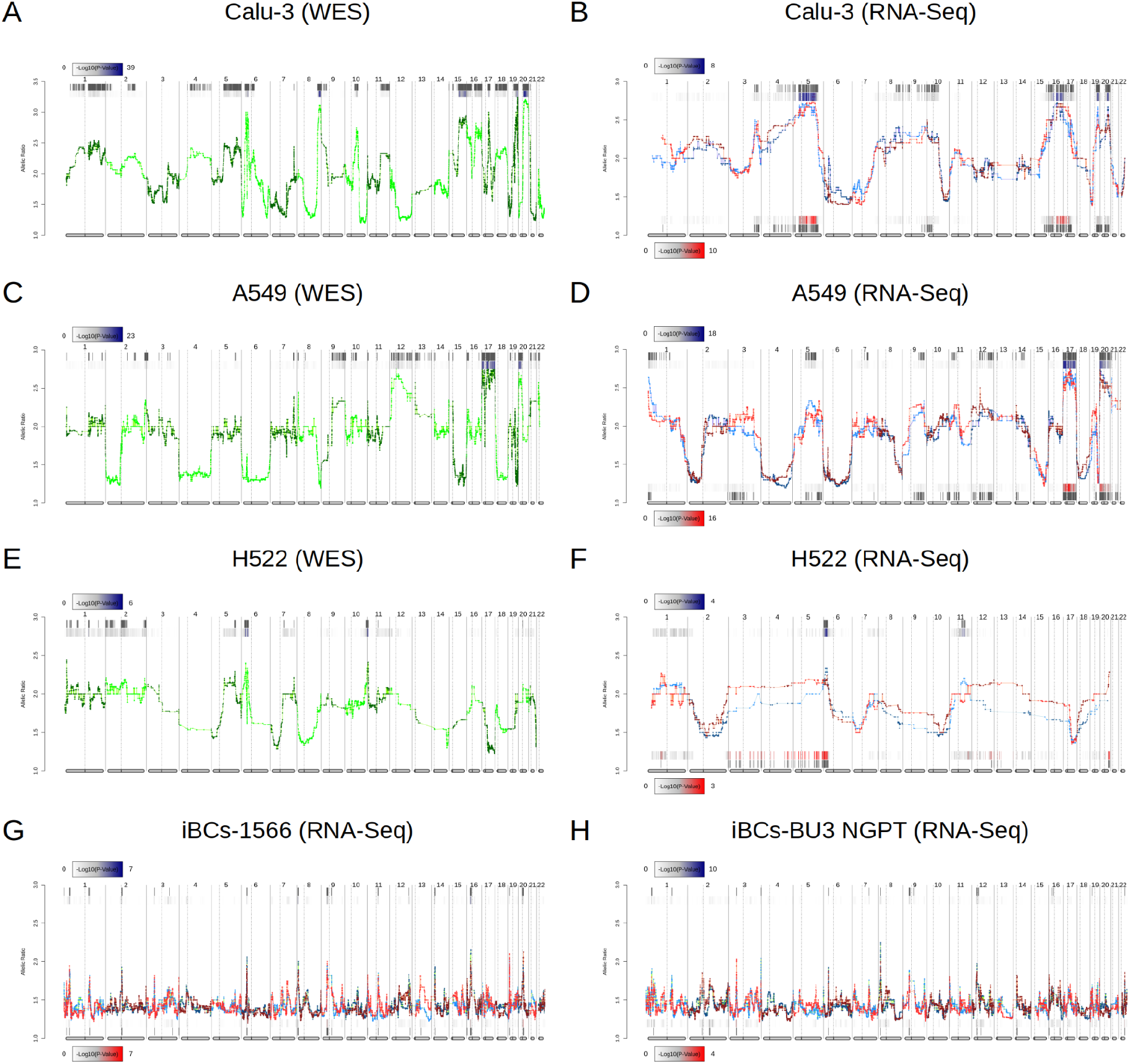
Comparison of chromosomal aberrations between mock-treated and SARS-CoV-2-infected cell lines. Comparison between e-Karyotyping analysis of samples from whole-exome sequencing and RNA-Seq data from Calu-3 (A-B), A549 (C-D), and H522 (E-F). G-H) e-Karyotyping analysis in RNA-Seq data from iPSC-derived airway epithelium basal cells (iBCs) from 1566 and BU3 NGPT cell lines. For each experiment, red dots and lines represent SARS-CoV-2-infected replicates, whereas blue dots and lines show mock-treated replicates. Diploid samples usually display an allelic ratio (y-axis) around 1.4 as previously shown (43,81). The gray background shown at the top and bottom of each plot shows regions that reach statistical significance for aneuploidy using the piecewise constant fit algorithm. The color gradient displayed next to each region represents the FDR-corrected P value for both comparisons.

### Allelic imbalance at DASE sites is partly linked to the differential expression of haplotype blocks

To understand the extension of the allelic imbalance across neighboring eSNVs, we expanded our screening around the DASE sites of each gene. In seven experiments, the *BRD2, HLA-C, MAT2A, RPSA, SUPT6H*, and *TNFRSF11A* genes each displayed only one eSNV. Also, even though they had multiple eSNVs, the LogASE values of co-localized variants in four genes (*EHD2, GFM2, GSPT1, and UMPS*) did not change. We then sought to evaluate if these variations overlapped single gene isoforms. By mapping each DASE site and its nearby eSNVs to the transcripts, we saw that all of the eSNVs were in areas where more than one isoform passed through. Thus, the possibility of isoform-specific allele expression was excluded from this set of genes. Finally, for six genes in 14 experiments, we observed DASE sites neighboring eSNVs affected after SARS-CoV-2 infection. For instance, *HLA-B* displayed many SNVs impacted close to the DASE sites in Calu-3 and A549 cell lines. Similar results were found for the *HAVCR1*, *NQO2*, and *UVSSA* genes.

The overwhelming occurrence of eSNVs at the MHC locus raises the question of whether the eSNVs are in phase, i.e., in the same RNA molecule and transcribed from the same parental allele. We use HapTree-X to reconstruct longer-range haplotypes using allelic imbalance at theoretically heterozygous eSNVs (44). We focused our analysis on six genes with DASE sites that span at least two heterozygous SNVs. We reconstructed the phased haplotype for all genes investigated across the different experiments. DASE sites affected by SARS-CoV-2 infection co-localized on the same RNA molecules raising the possibility of viral-induced differential haplotype expression (DHE) (Figure 4). This pattern was consistently observed in the *HLA-B* gene throughout seven different comparisons. DHE also occurred in the *HLA-C* and *UVSSA* genes in at least two comparisons.

**Figure 4.**
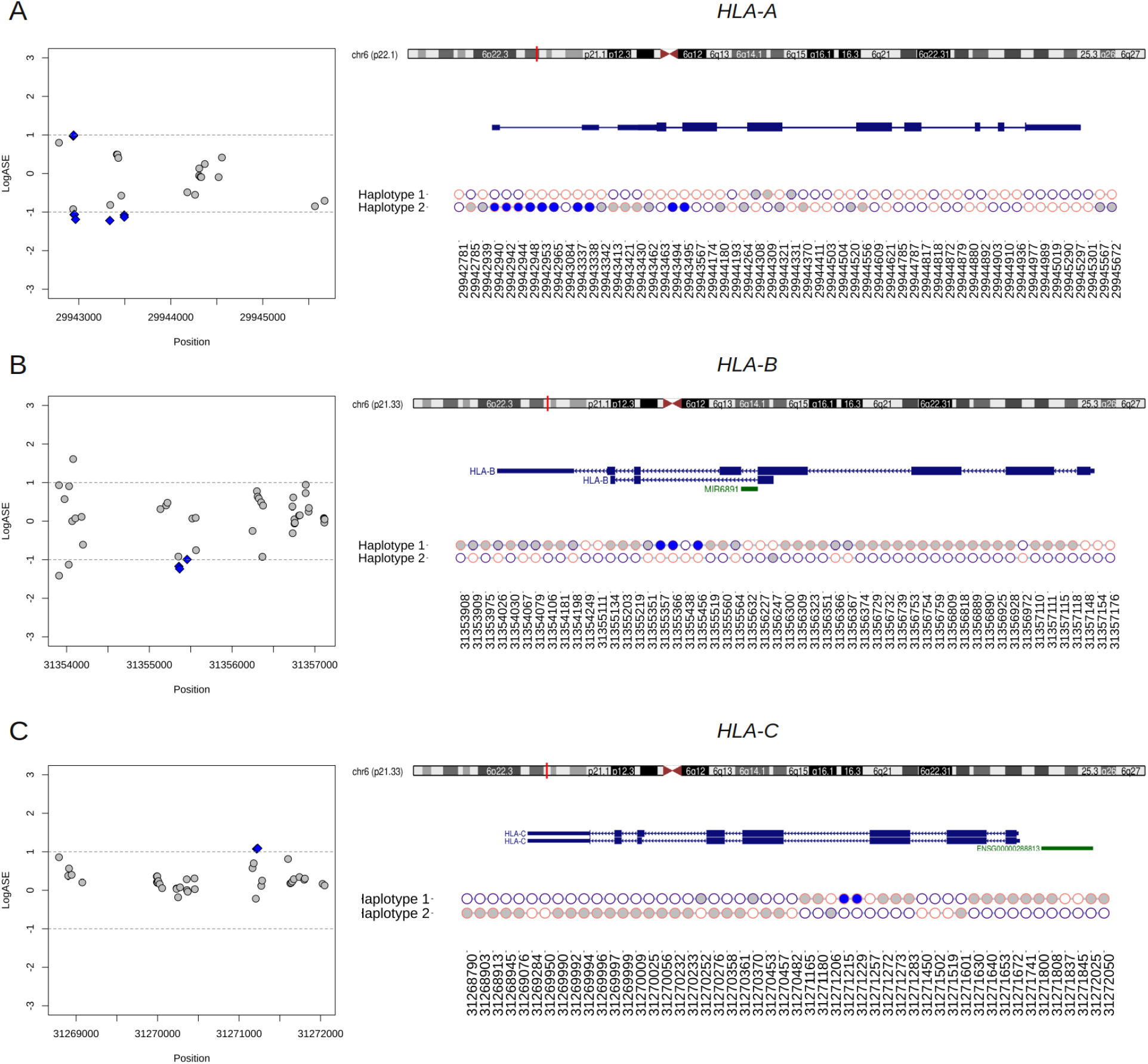
Phasing of DASE sites and co-localized eSNVs from classical MHC class I genes using RNA-seq data from Calu-3 cells. A-C) Regional plot of eSNVs localized around the *HLA-A, -B*, and *-C* genes. In blue, DASE sites revealed by the binomial test. The gray represents the other eSNVs tested that did not reach statistical significance. The ideogram of the chromosome is also shown, and a red tick shows where each relevant transcript isoform is located. Next, a plot showing the single haplotype block spanning the genes under analysis. Purple circles represent the reference allele, while the alternative is represented in pink. The x-axis refers to the genomic position of each eSNV in the GRCh38 genome assembly, and the y-axis shows the two haplotypes from the chromosomal locus. Blue and gray circles matched the SNVs in the regional plot on the left. The SNVs found in HLA genes that were not used in the binomial test are shown by the empty circles.

A single haplotype block spanned the entire *HLA-B* gene, covering a genomic window of 3,268 bp in Calu-3 cells (Figure 4). We identified 80 SNVs, of which 44 were interrogated during DASE analysis, five of which were differentially expressed. All the other affected eSNVs that did not reach statistical significance or pass the LogASE threshold were in phase with DASE sites. In the other experiments, the reconstruction of the HLA-B haplotype was fragmented, but we could identify haplotype blocks harboring DASE sites and co-localized eSNVs where all alleles with distinct expression patterns during viral infection were phased. Single haplotype blocks were also detected in *HLA-A* and *HLA-C* (Figure 4). Ten DASE sites were detected in *HLA-A* in Calu-3 and were co-expressed with the 18 eSNVs in Haplotype 2 (Figure 4). For *HLA-C*, we noticed DHE toward the haplotype #2, similar to that observed for non-HLA genes such as *HAVCR1* and *UVSSA*.

### DASE sites and affected co-localized SNVs discriminate MHC class I alleles preferentially expressed during infection

The reconstruction of extended haplotype blocks in the MHC class I locus allows allelic typing, which provides insights into the preferential expression of alleles during SARS-CoV-2 antigen presentation. Thus, to predict the HLA alleles assigned to each haplotype reconstructed in the previous analysis, we performed sequence-based HLA typing from RNA-Seq reads in each sample. We identified six samples heterozygous for HLA alleles that displayed DHE, in which the DASE sites and co-localized SNVs could discriminate the HLA allele preferentially expressed (Table 1). The *HLA-A* gene of Calu-3 cells was heterozygous for the A*24:02 and A*68:01 alleles. The 10 DASE sites with increased expression after infection mapped to the A*68:01 allele. In the *HLA-C* gene from Calu-3, the DASE sites did not distinguish the heterozygous HLA alleles. Using the SNVs in phase with DASE sites, we could differentiate the imbalance between the two alleles. We found the preferential expression of the allele C*15:02 co-expressed with the allele C*07:02 with three discriminant variations. The DASE sites found in *HLA-B* of Calu-3 did not distinguish alleles. However, by extending our analysis to three neighboring SNVs, we found an imbalance between the alleles B*51:01 and B*07:02. We observed that two altered SNVs mapped to B*51:01 while B*07:02 was characterized by a single variant (Table 1). Lastly, we found that the B*44:03 and B*18:01 alleles of the *HLA-B* gene were both heterozygous in the A459 experiments.

## Discussion

This study identified an imbalanced expression of genetic variations in classical MHC class I genes and ten other genes associated with SARS-CoV-2 infection. Gene ontology analysis showed that the 13 genes with DASE sites in Calu-3, A549, and H522 are enriched in protein binding functions, some of which are involved in SARS-CoV-2 infection, COVID-19 disease progression, and severity. *HLA-A*, *-B*, and *-C* genes act on endogenous peptide antigen presentation and are associated with disease susceptibility. The transcriptional regulator bromodomain-containing protein 2 (BRD2) is a potent regulator of ACE2 transcription in Calu-3 cells (48). The EHD2 protein, highly enriched at the neck of caveolae, controls a cell-autonomous, caveolae-dependent fatty acid uptake pathway by adipocytes, endothelial cells, and muscle cells (49). Importantly, *EHD2* is underexpressed in obese patients, a known risk comorbidity for severe COVID-19.

Patients carrying the cytosolic glutathione S-transferase *GSPT1* rs1695 allele are at lower risk of COVID-19 development (50). The hepatitis A virus cellular receptor (HAVCR1, also called KIM1), used by Ebola, Marburg, Dengue, and Zika viruses, is an entry factor for SARS-CoV-2 to kidney cells, where the virus induces organ abnormalities associated with poor prognosis and mortality in COVID-19 patients (51). The methionine adenosyltransferase 2A (*MAT2A*), involved in S-adenosylmethionine methylation pathways, is differentially upregulated in mono-CD14+CD16+ cells in patients with severe COVID-19 (52). MAT2A presumably is required to methylate the SARS-CoV-2 RNA cap structures, allowing genome transcription and preventing the recognition of RNA Cap structures by cellular innate immunity receptors (53). The *SUPT6H* gene codes one of the many RNA binding proteins profoundly down-regulated upon SARS-CoV-2 infection (54). The uridine monophosphate synthase (UMPS) is involved in pyrimidine biosynthesis, and pyrimidine inhibitors synergize with nucleoside analogs to block SARS-CoV-2 replication (55).

The observed allele bias in classical MHC class I genes leads to the preferential expression of one allele within a heterozygous locus, showing that the upregulation of these genes is driven in a haplotype-specific manner. The classical MHC class I molecules handle mainly self-peptides or viral antigens. The exposure of the HLA-peptide complex on the cell surface is followed by CD8+ cytotoxic T lymphocyte binding, which may induce apoptosis in virally infected cells and generate long-term immunological memory. By having heterozygous alleles in the *HLA-A*, *-B*, and *-C* genes, up to six MHC class I alleles can be expressed at a time in a single human cell. Perturbations in MHC allelic expression can change how antigens are presented. The cellular immunity conferred by CD8+ memory T cells is crucial to fighting earlier SARS-CoV-1 infection and the current SARS-CoV-2 pandemic, even with or without humoral responses (56–59). Though our findings seem to be limited to the repertoire of antigens for T CD8+ cell presentation, the isoform expressed may play a role in the efficiency of the immune response to viral infection.

For example, the A*68:01 allele overexpressed in Calu-3 has been predicted to have a high binding affinity to SARS-CoV-2 epitopes (60). A*68:01 is a common allele found across different populations at a frequency of 5.2–25%. This allele was strongly associated with mortality from influenza A (H1N1) infection (61,62). A large-scale analysis also revealed a proclivity for the worst COVD-19 outcome in patients with the B*51:01 allele that is overexpressed in Calu-3-cells (63). *In silico* analysis identified a high affinity for potential T-cell epitopes of S-protein (64). Previous studies reported a protective role of B*51:01 in the long-term control of AIDS progression in HIV-infected individuals (65–67). The alternative allele B*07:02, co-expressed with B*51:01, had a beneficial association with high antiviral efficacy against SARS-CoV-2 (68). For *HLA-B* of Calu-3 cells, we were not able to determine the phase of DASE sites considering the two alleles B*51:01 and B*07:02.

Cross-referencing with the HLA peptidome in Calu-3 infected by SARS-CoV-2 revealed that the epitopes presented on the cell surface matched most of the HLA alleles that were found to be differentially expressed by our analysis (69). The majority of peptides presented by HLA-A on the Calu-3 surface matched the A*68:01 allele. Nagler and his team did not see B*51:01 and C*07:02 being expressed in SARS-CoV-2-infected Calu-3. These results may help settle the disagreement about the haplotype that is most strongly expressed at the RNA level for *HLA-B*. Thus, the absence of epitopes matching the alternative allele for both HLA-*B* and *HLA-C* genes shows that the differential haplotypic expression may be reflected at the protein level. It is still not clear if the immunodominant epitope controls the preferential expression of the HLA alleles or if the different expression of the HLA alleles makes some peptides more likely to be chosen. The three HLA alleles upregulated in Calu-3 may play a protective role against COVID-19 (Figure 5). A*68:01 showed a protective effect against severe manifestations of the disease in Tapachula-Chiapas, Mexico (70). In contrast, the peptides presented by A*68:01 derived from the envelope protein are homologous to the neuronal cell adhesion molecule (NCAM) (71). Thus, A*68 has been associated with developing Guillain-Barre syndrome (GBS). B*07:02 and C*15:02 have antiviral activity and resistance against SARS-CoV-2, respectively (68,72).

**Figure 5.**
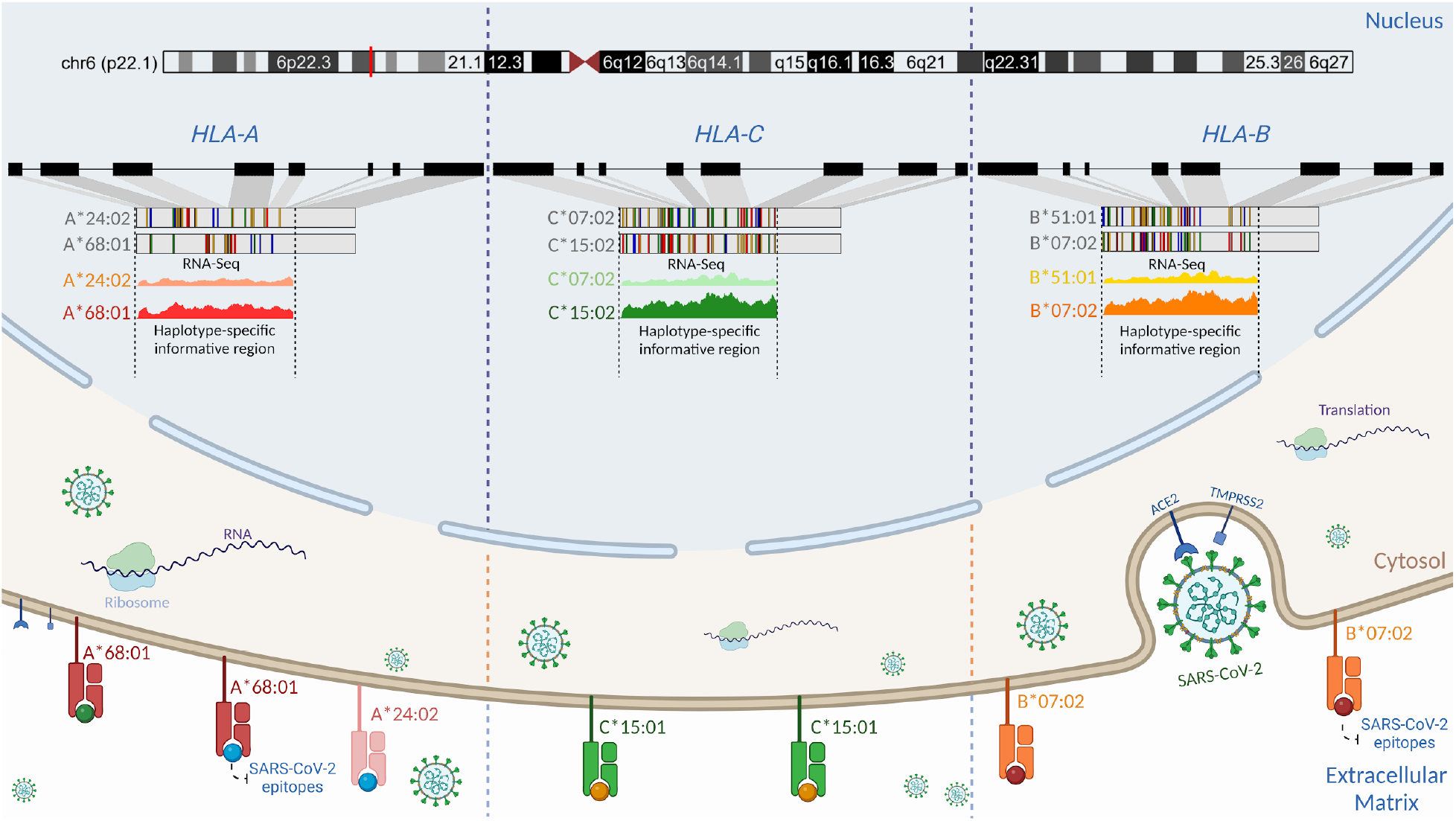
Schematic representation of the proposed regulatory genetic mechanism associated with the haplotype-specific expression of class I HLA alleles during SARS-CoV-2 infection. Viral cell entry triggers preferential transcription of the RNA molecules in the classical class I HLA locus. Even though Calu-3 is heterozygous for *HLA-A, -B*, and *-C* alleles (69), we found that the A*68:01 allele was more expressed than the A*24:02 allele, the B*07:02 allele was more expressed than the B*51:01 allele, and the C*15:02 allele was more expressed than the C*07:02 allele. Such differences in the expression may be attributed to structural differences in promoter motifs (82), transcriptional factors, genetic variations, and environment (83,84). Cross-referencing analysis using HLA peptidome data from Calu-3 infected by SARS-CoV-2 generated by Nagler and colleagues (2021) revealed that the same RNA molecule found to be preferentially expressed in RNA-Seq data corresponds to the HLA protein expressed on the cell surface for classical class I alleles. Nagler and colleagues (2021) reported that most peptides presented on the cell surface matched the A*68:01 allele when compared to the A*24:02. For *HLA-C*, no peptide matching C*07:02 was found after infection. Similarly, the B*07:02 allele was found to be preferentially expressed at the HLA-B locus. By using DASE, swapping alleles with low binding affinity could be a part of the defense that helps COVID-19 outcomes be less severe. Created with BioRender.com.

In the *HLA-B* gene in A549 cells, we observed the preferential expression of the B*18:01 allele over the B*44:03 allele in all experiments. B*18:01 was associated with the manifestation of subacute thyroiditis triggered during the SARS-CoV-2 infection (73). This allele has also been linked to T cell cross-reactivity between EBV epitopes and a self-peptide, causing an aberrant immune response (74). The HIV viral replicative capacity was significantly higher in subjects expressing the B*18:01 allele (75). In contrast to the patterns seen with *HLA-A* and *-B*, there is no clear link between the allele C*15:02 and COVID-19. This allele was upregulated in Calu-3 experiments when co-expressed with C*07:02. The C*15:02 allele confers resistance against SARS-CoV infection (72). Francis and colleagues recently described the HLA-B*07:02 allele as presenting homologous epitopes from SARS-CoV-2 and other HCoVs, providing high pre-existing immunity. Preferential expression of HLA alleles may be closely connected to TCR repertoire diversity (76). Moreover, HLA genotypes and CD8+ T cell responses have been described as having implications for herd immunity and strategies to consider during vaccine design to guarantee long-term immunity against SARS-CoV-2 (76,77).

Zhang et al. reported allelic imbalances across HLA-B alleles in lung cell lines infected by SARS-CoV-2 using an alternative methodological approach. The authors offered three non-exclusive biologically plausible mechanisms to explain the differential haplotype expression: (i) the activation/silencing of one allele is attributed to pathological effects, (ii) independent regulation of the transcription of both alleles, and (iii) the presence of cis-acting regulatory elements (27). In our study, the occurrence of DASE sites in the *BRD2* gene mapping to the HLA chromosomal region corroborates the cis-acting regulatory elements’ hypothesis. During T cell activation, allele-specific expression changes were described in HLA and other autoimmune loci for CD4+ T cells (78).

ASE perturbations are not mechanistically unique to SARS-CoV-2 infection, despite the reported shift in allele expression in HLA and ten other genes. Multiple ASE alterations have been identified in CD4-T cells infected by the oncogenic Marek’s Disease herpesvirus (MDV) (79). MDV caused ASE changes in six genetic resistance loci (*MCL1, SLC43A2, PDE3B, ADAM33, BLB1*, and *DMB2*) that are related to T-cell activation, T-cell and B-cell receptors, ERK/MAPK, and PI3K/AKT-mTOR signaling pathways, all of which play important roles in MDV infection. Because ASE-affected genes represent the complex trait of genetic resistance to Marek’s disease, the trait is then determined by transcriptional regulation (80). Our results show that when SARS-CoV-2 infects cells, there is a transcriptional allelic flip in the affected genes, which occurs regardless of compensation of gene expression. We hypothesize that when the virus enters the cell, a DASE flip regulatory mechanism swaps HLA alleles that display epitopes with poor binding affinity. Functional studies are required to assess the biological significance of the transcriptional allelic flip. We warn against drawing any more conclusions from these results because the sample size was small and the study was done with public secondary WES and RNA-Seq data.

## Data Availability Statement

All the datasets generated for this study are included in the article and supplementary material. The NGS data (RNA-Seq and WES) used in this manuscript are from publicly available BioProjects. The primary data can be accessed from the NCBI SRA repository (https://www.ncbi.nlm.nih.gov/sra) under the ID listed in Table S1.

## Acknowledgments

We are very grateful to the authors and institutions that made the RNA-Seq and WES data publicly available. This work was developed in the framework of Corona-ômica-RJ (FAPERJ = E-26/210.179/2020 and E-26/211.107/2021); FAPERJ E-26/210.681/2021. ATRV is supported by CNPq (307145/2021-2) and FAPERJ (E-26/201.046/2022). EMA is supported by CNPq (308955/2019-6). RSFJ was a recipient of a graduate fellowship from CNPq. We gratefully acknowledge the assistance of the Rede Corona-ômica BR MCTI/FINEP (FINEP 01.20.0029.000462/20, CNPq 404096/2020-4).

## Statement of contribution to the field

In response to infection with SARS-CoV-2, cells engage their innate and adaptive immune systems. SARS-CoV-2 interferes with cytokine signaling and affects the antigen-presenting function of HLA molecules because of transcriptional modifications influenced by interindividual variance. By evaluating allele-specific expression as a measure of interindividual variation, we found that infection of human epithelial lung and airway cell lines with SARS-CoV-2 causes a shift in the expression of HLA alleles that swaps alleles with poor epitope binding affinity. The observed shift in HLA allele expression may relate to improved COVID-19 outcomes.

## Table Legends

Table 1. Detection of DASE sites and affected co-localized SNVs in MHC class I alleles.

Table S1. Public RNA-Seq and WES experiments included in this study.

Table S2. DASE sites identified in Calu-3, A549, and H522 cell lines.

Table S3. DASE sites found in airway epithelium basal cells derived from iPSCs.

